# Synthetic germ granules reveal a direct role of Vasa/DDX4 in RNA localization and translational activation

**DOI:** 10.64898/2026.02.01.703065

**Authors:** Ruoyu Chen, Henoc Zinga, Jay S. Goodman, Ruth Lehmann

## Abstract

Cytoplasmic RNA granules, including stress granules, P bodies, neuronal RNA granules, and germ granules, are essential for RNA storage and regulation across a wide range of organisms. However, dissecting the contributions of individual factors to granule function is challenging because of the interdependence of components *in vivo*. This is especially true for DEAD-box helicases, common regulators of mRNA granules, whose specific contributions remain unclear. In this study, we developed a synthetic approach to de novo generate germ granules, enabling us to identify the minimal machinery needed for RNA localization and translational activation. Using a self-assembling PopTag-based scaffold derived from Caulobacter fused to the RNA-binding domain (RBD) of the germplasm organizer Oskar, we found that the recruitment of endogenous germ granule mRNAs (*nanos* and *pgc*) depended on the DDX4 protein Vasa. By employing orthogonal RNA tethering approaches, we demonstrate that Vasa is both necessary and sufficient for localized mRNA translation. Consistent with these findings, acute depletion of Vasa from endogenous germ granules specifically reduced Nanos translation without affecting mRNA localization, confirming Vasa as a core factor linking RNA recruitment to localized translational activation. These *in vivo* reconstitution experiments reveal a two-component module in which a scaffold RBD and the Vasa helicase, but not other DEAD-box helicases, enable RNP condensates to accumulate specific RNAs and promote their translation. Overall, our study uncovers previously unrecognized functions of an RNA helicase within ribonucleoprotein condensates and demonstrates the power of synthetic biology to analyze complex biomolecular condensates in living organisms.

## Introduction

Ribonucleoprotein (RNP) granules are prevalent, membrane-less cellular structures that enrich a wide variety of RNA and RNA-binding proteins (RBPs), serving as a central hub for post-transcriptional regulation of gene expression. Classic and best-characterized examples include stress granules, processing bodies, and nucleoli^1,2^. DEAD-box RNA helicases, a large class of ATPases that unwind double-stranded RNA, have been found in virtually all types of RNP granules, like eIF4A and DDX3 in stress granules, DDX6 in processing bodies, and DDX21 in nucleoli ^3^. Due to their ability to disrupt RNA-RNA interaction, the activation of DEAD-box helicase has been associated with the dissolution and turnover of RNP granules ^4,5^. Conversely, inactive helicases are thought to drive RNP granule assembly, mediated by the intrinsically disordered regions (IDRs) of the helicases and stabilized RNA-RNA interactions ^4^. It remains unclear whether and how RNA helicase activity can positively influence RNP granule assembly, and whether RNA helicases can be directly involved in mRNA translation within an RNP granule.

Germ granules are a specialized type of RNP granules that are crucial for germline and embryonic development^6,7^. In several species, such as fruit flies, zebrafish, and frogs, germ granules form in a specific region of the oocyte called the germ plasm, from which primordial germ cells will develop during embryogenesis. These granules are essential for stabilizing and translating specific mRNAs, including those that determine germ cell fate^6–8^. Vasa/DDX4 is a conserved DEAD-box helicase expressed in the germ cells of all animals^9–14^. In germ cells, Vasa is a major component of germ granules and serves as a universal germ granule marker ^10^. Nevertheless, the precise role of Vasa in germ granules remains unclear. For instance, in Drosophila, it is uncertain whether Vasa has a direct role in assembling germ granules, since Vasa is also necessary for producing and stabilizing Oskar, the scaffold protein essential for germ granule formation^15–19^. It is also not known whether Vasa directly controls the translation of germ granule-associated mRNA, as none of the *vasa* null or loss-of-function mutant embryos form germ granules^20,21^.

In addition to Vasa and Oskar, germ granules contain a complex suite of proteins, including the evolutionarily conserved germline proteins Tudor and Aubergine^22,23^. Many of these components have been implicated in the assembly and/or the downstream function of germ granules. However, the minimal requirements for building a functional germ granule remain undefined, leaving the principles of germ granule assembly and the mechanisms of RNA regulation within germ granules unclear. Assigning the function of each component using genetic characterization of mutant alleles has often been confounded by epistasis effects. For example, Aubergine plays a role in Oskar translation, which precedes germplasm assembly^24^. A bottom-up approach is required to confidently establish the role of each component.

Here, we used a novel *in vivo* reconstitution strategy to identify the minimal components required for the basic functions of germ granules in Drosophila. Specifically, we designed a synthetic scaffold protein that can self-assemble into germ granule-like condensates, localize specific germplasm mRNA, and activate localized translation in Drosophila embryos. By further dissecting the designer granules, we uncovered an essential role of Vasa in mediating RNA localization. Combined with quantitative imaging of translation *in vivo*, we directly demonstrate the necessity and sufficiency of Vasa in activating the translation of granule-localized mRNA. Specifically, Vasa utilizes its RNA unwinding activity to protect granule-localized *nanos* mRNA from Smaug-mediated repression. Despite being indispensable for germ granule assembly, Vasa can be specifically depleted after assembly without affecting other granule components. Targeted depletion of Vasa in native germ granules confirms our reconstitution-based results.

## Result

### *In vivo* reconstitution of functional RNP granules using a designer construct

To define the minimal components required for germ granules to mediate RNA localization and translational activation, we sought to functionally reconstitute germ granules in Drosophila embryos using an artificially designed granule-forming protein. Like most biomolecular condensates, germ granule formation is driven by a scaffold protein possessing both self-assembly and RNA-binding properties, exemplified by Oskar in *Drosophila*, Xvelo in *Xenopus*, and Bucky Ball in *Danio*^6–8,25,26^. While the scaffold proteins are highly divergent, Vasa is a conserved component of germ granules across species. In *Drosophila*, Vasa is recruited to germ granules via direct interaction with the N-terminal LOTUS domain of Oskar^27,28^. We hypothesized that a synthetic protein combining self-assembly, RNA-binding, and Vasa-recruitment properties could form designer condensates that recapitulate the functions of germ granules. To this end, we employed NanoPop—a modular construct consisting of an anti-GFP nanobody fused to PopTag, an oligomerization domain derived from the bacterial protein PopZ^29^ (Figure 1A). Within the NanoPop construct, we inserted the RNA-binding domain of Oskar (OSK domain) to confer RNA-binding capability and specificity^28,30,31^. The resulting construct, termed NanoPop::OSK, is expressed at the anterior pole of Drosophila embryos using a *bicoid (bcd)* 3’UTR to spatially restrict the granule formation^18,32^. VasaGFP is co-expressed in the embryos to bind NanoPop::OSK and form granules collectively at the anterior pole (Figure 1B, see Method for the details about NanoPop design and expression strategy).

**Figure 1.**
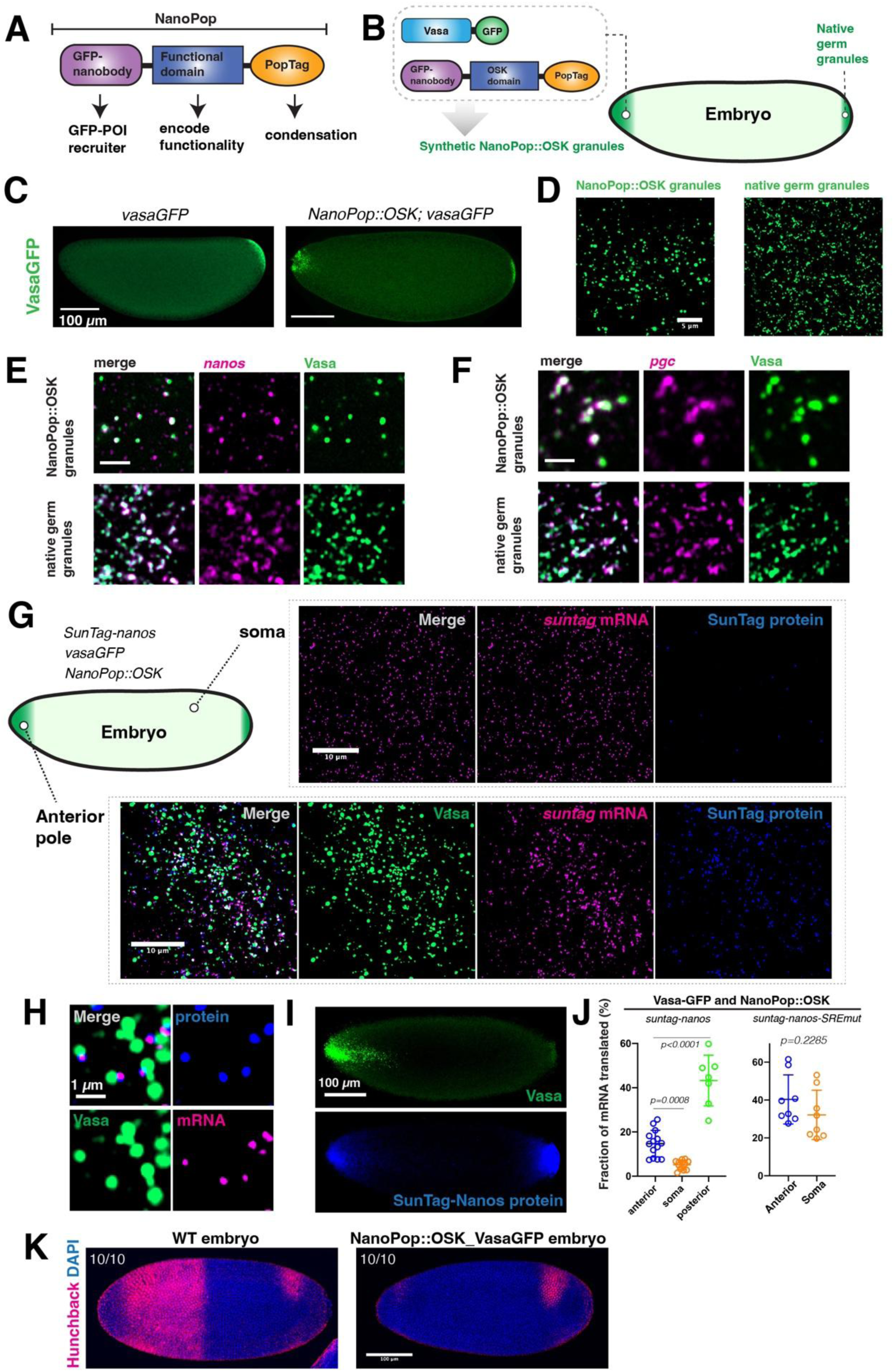
A. A schematic of NanoPop construct. B. Expression of NanoPop::OSK with VasaGFP in a Drosophila embryo. OSK domain of Oskar protein is used as the functional domain in the NanoPop construct. NanoPop::OSK is expressed at the anterior of the embryo. The anterior of embryos faces left in all figures. C. Left: an embryo expressing VasaGFP, which is enriched at the posterior germplasm. Right: an embryo expressing NanoPop::OSK-*bcd*3’UTR and VasaGFP. VasaGFP is enriched both at posterior and in the synthetic granules at the anterior. D. A close-up view of NanoPop::OSK granules and native germ granules labeled by VasaGFP. E. Localization of *nanos* mRNA (smFISH, magenta) to NanoPop::OSK granules and native germ granules labeled by VasaGFP (green). F. Localization of *pgc* mRNA (smFISH, magenta) to NanoPop::OSK granules and native germ granules labeled by VasaGFP (green). G. Translational activation of *suntag-nanos* mRNA by NanoPop::OSK granules (green). Translation is visualized by the colocalization of SunTag protein spots (anti-GCN4 staining, blue) with the *suntag* mRNA spots (smFISH, magenta). Translation in soma, which is largely repressed, is shown for comparison. H. A close-up view of translating mRNAs on NanoPop::OSK granules. I. Strong SunTag-Nanos protein staining at both the anterior and posterior poles of an embryo due to translational activation. J. Quantification of the percentage of mRNA being translated. K. Expression of Hunchback (anti-Hb, magenta) in WT and NanoPop::OSK_VasaGFP embryos. Nuclei were stained with DAPI (blue). All (10 out of 10) WT embryos showed Hunchback at the anterior, while all (10 out of 10) NanoPop::OSK_VasaGFP embryos showed no Hunchback at the anterior.

GFP-positive granules were formed at the anterior pole in the *NanoPop::OSK-bcd 3’UTR* and VasaGFP-expressing embryos (Figure 1C and D). Synthetic NanoPop::OSK granules resembled native germ granules in morphology (Fig. 1D and Fig. S1A, B). NanoPop::OSK granules tended to be larger and rounder while native germ granules often adopted elongated shape (Fig. 1D and Fig. S1B). Like endogenous germ granules, the synthetic granules often formed hollow spheres^33,34^ (Fig. S1A). Notably, *nanos* and *pgc* mRNA-- normally localized to germ granules at the embryo posterior^35^-- was also localized to the NanoPop::OSK granules, suggesting a specific mRNA binding property of the designer granule (Figure 1E, F). In germ granules, localized *nanos* mRNA is actively translated, while unlocalized *nanos* is translationally repressed mainly by the translational repressor Smaug^33,36–40^. Localized translation of nanos produces a Nanos protein morphogen gradient that determines the posterior fates of the embryo, and high levels of Nanos protein are required for germ cell specification. To ask whether *nanos* mRNA localized to NanoPop::OSK granules was actively translated, a *suntag-nanos* transgene was expressed to visualize the translation of *nanos* at the single-molecule level, together with *NanoPop::OSK-bcd 3’UTR* and VasaGFP^33,40^. Localized *suntag-nanos* mRNA in the NanoPop::OSK granules showed significantly higher translation compared to the unlocalized mRNA in the soma (Figure 1G-J). The mRNA of *Suntag-nanos-SREmut*, bearing a mutated Smaug response element (SRE) and thus not repressed by Smaug, displayed uniformly elevated translation both in soma and in NanoPop::OSK granules (Fig. 1J, Fig S1C). This suggests that NanoPop::OSK granules activate translation by lifting Smaug-mediated repression, a mechanism similar to native germ granules. It should be noted that, however, translation activation by the NanoPop::OSK granules was not as strong as the activation conferred by native germ granules or by SRE mutation, suggesting a compromised function of the designer granule (Figure 1J). Nevertheless, NanoPop::OSK granules created sufficient Nanos protein at the anterior to fully inhibit the translation of the anterior determinant Hunchback, recapitulating the morphogenic function of native germ granules (Fig. 1K)^41^.

While capable of localizing *nanos* RNA and generating a Nanos protein gradient sufficient to suppress hunchback translation, synthetic granules did not replicate all the functions of germ granules. Anterior expression of *oskar* can lead to functional ectopic germ granules, including the induction of anterior pole cells, the primordial germ cell precursors^18^. However, pole cells did not form at the anterior of embryos expressing NanoPop::OSK-bcd3’UTR and VasaGFP, indicating that the synthetic granules fail to induce factors needed for pole cell formation (Fig S1D and E). Consistent with this result, mRNA of *gcl*, an essential gene for pole cell formation^42,43^, showed only weak localization to the synthetic granules (Fig. S1F). Together, this showed that NanoPop::Osk granules exhibited partial functional overlap with native germ granules. What other factors are needed to recruit the remaining suite of essential germplasm mRNAs will be the focus for a future study.

### Vasa is required for RNA localization to germ granules

The modular design and programmability of the NanoPop system enable functional dissection of granule components. We focused first on the ability of NanoPop granules to localize *nanos* RNA. To ask how the OSK RNA-binding domain contributes to granule assembly and RNA localization, we generated a NanoPop construct lacking the OSK domain (NanoPop-*bcd*3’UTR) and co-expressed it with VasaGFP (Fig. 2A). The resulting NanoPop_VasaGFP granules could not localize *nanos* mRNA, confirming the role of the OSK domain in *nanos* mRNA localization. Moreover, while NanoPop was expressed at the anterior using *bcd* 3’UTR, NanoPop_VasaGFP granules were not restricted to the anterior pole but rather distributed throughout the embryos, suggesting that the OSK RNA-binding domain and its ability to recruit mRNA are critical for restricting the mobility of granules within embryos (Fig. 2A). To explore whether the OSK domain determines the specificity of RNA localization to germ granules, as indicated by the high enrichment of certain RNAs, such as *nanos*, in germ granules ^35,44^, or whether it simply offers a broader affinity for RNA aiding in granule formation, as suggested by *in vitro* experiments that showed only low sequence specificity of the OSK domain for RNA^30^, we used the NanoPop system to test other RNA binding domains (RBD). We inserted the RBD of human G3BP1, an RNA-binding protein known to bind mRNA with low specificity, into the NanoPop^45^. The resulting NanoPop::G3BP1 formed granules with VasaGFP that remained confined to the anterior but failed to localize nanos mRNA (Fig 2A, B), suggesting that the OSK domain encodes sequence specificity for targeting certain mRNAs to germ granules, while RNA recruitment itself may help stabilize the synthetic granules.

**Figure 2.**
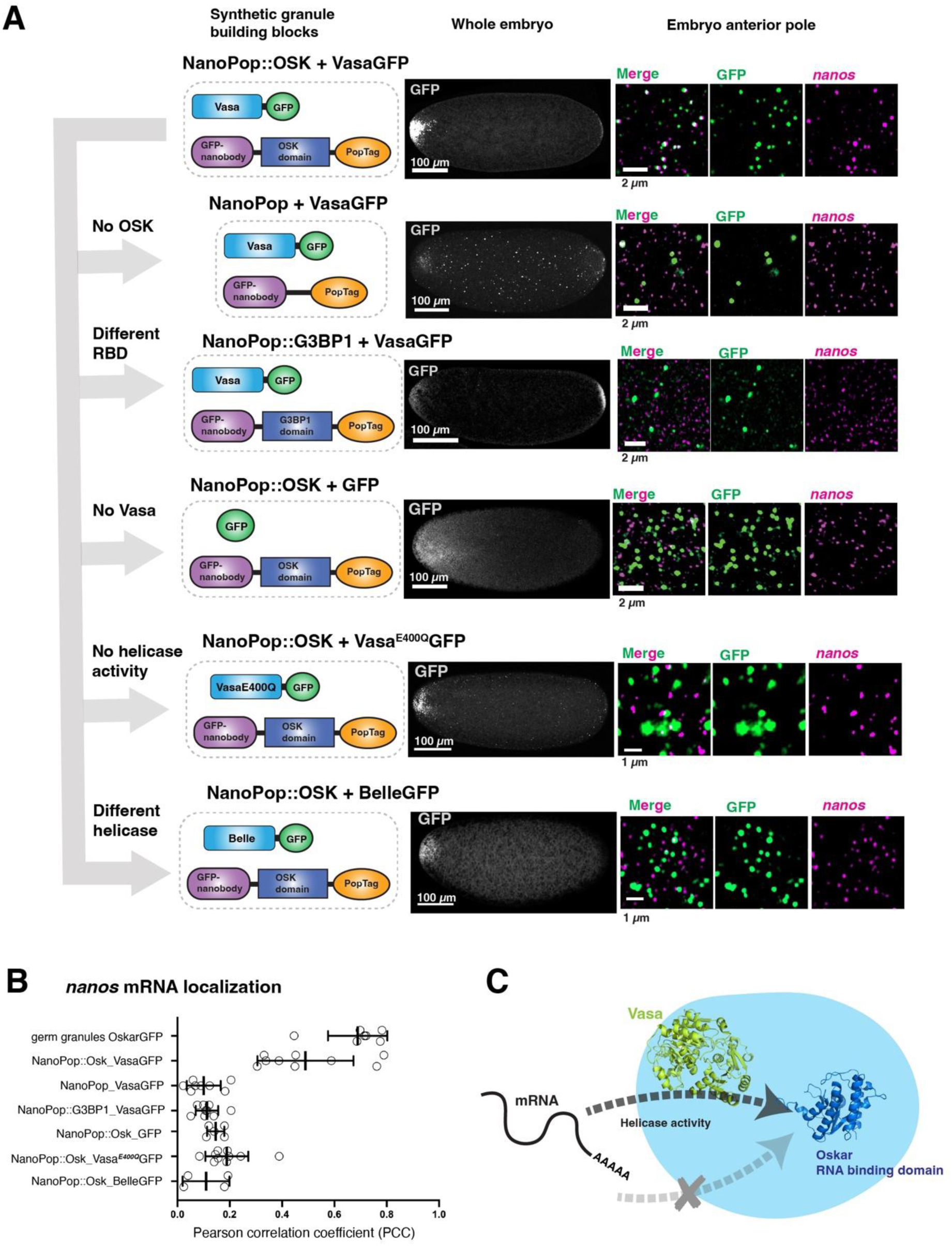
A. Different synthetic NanoPop granules and assessment of *nanos* localization. Indicated NanoPop constructs were expressed at the anterior pole using *bcd* 3’UTR. GFP or GFP fusion proteins was co-expressed. An example image of embryo is shown on the left. On the right shows a zoom-in of the anterior with NanoPop granules, labeled by GFP (green), and *nanos* mRNA labeled by smFISH (magenta). B. Quantification of the Pearson Correlation Coefficient (PCC) between *nanos* RNA and NanoPop granules as a measurement of RNA localization. C. A hypothetical model of *nanos* localization to germ granules. *Nanos* does not localize by directly binding to Oskar. Instead, *nanos* first interacts with Vasa, which pass *nanos* onto Oskar using helicase activity.

Our results suggest that the OSK domain provides RNA localization specificity. Thus, in theory, combining NanoPop’s ability to create a condensate scaffold with the OSK domain’s capacity to bind RNA should be sufficient to recruit mRNA. We found, however, that granules made by NanoPop::OSK with GFP instead of VasaGFP were not able to localize *nanos*, suggesting that Vasa, in addition to Oskar, is indispensable for RNA localization to the designer granules. An E-to-Q mutation in the conserved DEAD motif in the ATPase domain (E400Q) disables the helicase activity of Vasa locking the helicase in a closed, RNA-bound conformation^46^. Granules made by Vasa^E400Q^GFP and NanoPop::OSK failed to localize *nanos*, suggesting that RNA unwinding by Vasa is involved in RNA localization to the designer granules. Moreover, the evolutionarily most related paralog of Vasa, Belle (DDX3), could not replace Vasa for RNA localization even though its GFP fusion proteins could form granules with NanoPop::OSK, suggesting Vasa has a unique function to mediate RNA localization to designer granules (Fig. 2A, B).

The results above suggested that Vasa is needed for RNA to bind the RBD of Oskar and become localized to germ granules. This function has been unappreciated so far, because Vasa has been implicated in Oskar protein synthesis and accumulation during oogenesis, which preceded this later role in RNA localization within granules^16,19^. Indeed, in *vasa* null mutant embryos, Oskar appeared weak and diffuse at the posterior, and germ granules rarely formed, preventing us from assessing Vasa’s role in RNA localization (Fig S2A). We found that in the embryos of two *vasa* mutants, I256N and R378A, both of which impair RNA binding^20,21^, Oskar was able to accumulate and form aggregates at the posterior but *nanos* mRNA was not localized to these aggregates (Fig. S2A). This result supports our conclusion that RNA-Vasa interaction is required for RNA localization and Oskar alone is not sufficient (Fig. 2C).

The Piwi family protein Aubergine (Aub) and its bound piwi-interacting RNAs (piRNAs) are one of the major components of germ granules and have been implicated in recruiting mRNA via base-pairing between piRNA and mRNA^23^. We asked whether Aub was involved in *nanos* mRNA localization to the designer NanoPop::OSK_VasaGFP granules. Immunofluorescence showed that, unlike native germ granules, which were highly enriched with Aub, Aub was not localized to the designer granules, suggesting that Aub is not necessary for *nanos* localization (Fig. S2C). A designer granule formed by NanoPop and AubGFP did not localize *nanos* mRNA, suggesting that Aub alone is not sufficient to recruit specific mRNA to granules (Fig. S2D). Together, these data show an indispensable role of Vasa in RNA localization to designer granules and germ granules. Our data align with a model in which Vasa facilitates RNA localization by using its RNA-unwinding activity to transfer mRNA to the RNA-binding protein in the RNP granules, which is Oskar in this case.

### Vasa is necessary and sufficient to activate translation in reconstituted germ granules

Vasa/DDX4 has been implicated in activating mRNA translation in various species, but the evidence has been correlative^47–50^. In Drosophila, it remains unclear if Vasa directly affects the translation of germ granule-localized mRNAs. Addressing this question is particularly challenging, as Vasa is required for germ granule assembly and initial RNA recruitment to the granule. We bypassed the need for Vasa to localize RNA by constructing a designer granule with the MS2-MCP system (Fig 3AB). MS2 is a stem-loop RNA sequence derived from MS2 bacteriophage and is bound by MS2 coat protein (MCP) with high specificity and affinity^51^. We generated a transgene carrying *suntag-nanos* with 24 copies of MS2 sequences inserted into the 3’ UTR (*suntag-nanos-MS2×24*). A designer granule formed by NanoPop and MCP-GFP could localize the *suntag-nanos-MS2×24* mRNA independent of Oskar or Vasa. (Fig. 3A and Fig. S3A). Localized suntag-nanos-MS2×24 mRNA remained translationally repressed (on average, less than 10% of localized mRNA was translationally active), suggesting that localization to a granule or simply increasing concentration is not enough to activate translation^51^, and that both NanoPop and MCP-GFP have little to no effect on mRNA translation (Fig. 3B-D). Embryos expressing NanoPop, MCP-GFP, and Vasa-GFP generated designer granules with both MCP-GFP and Vasa-GFP (Fig. 3A and Fig. S3B). Remarkably, *suntag-nanos-MS2×24* mRNA localized to the NanoPop_MCP-GFP_Vasa-GFP granules showed significantly increased translation (on average, 30% translationally active) compared to the NanoPop_MCP-GFP granules. Thus, Vasa alone was sufficient to activate translation when being targeted to a designer mRNA granule (Fig. 3B-D). *Suntag-nanos* mRNA lacking MS2 sequences did not localize to the NanoPop_MCP-GFP_Vasa-GFP granule (Fig. S3A and C). Neither did it increase translation at the anterior of the embryo due to the presence of these granules, suggesting that Vasa specifically activated translation of mRNA in its vicinity (Fig. S3C and D).

**Figure 3.**
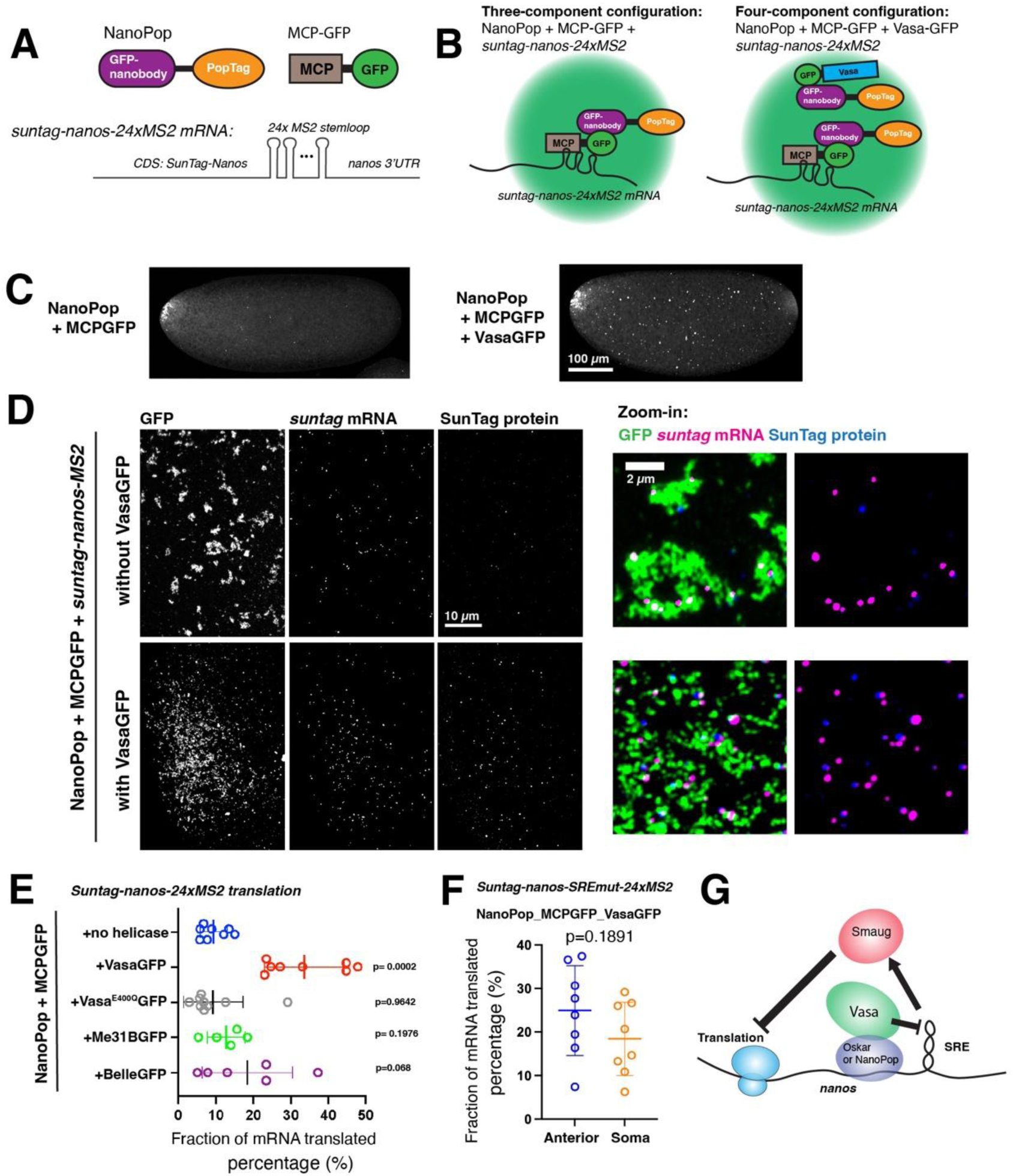
A. Components for building a synthetic granule with constitutive RNA localization. B. Schematics of hypothetical synthetic granules. Left: synthetic granules made of NanoPop and MCP-GFP. *Suntag-nanos-24xMS2* mRNA localizes to the granule through MS2-MCP interaction. Right: both MCP-GFP and Vasa-GFP should co-assemble with NanoPop into one synthetic granule. C. Left: an embryo expressing NanoPop-*bcd*3’UTR and MCP-GFP. Right: an embryo expressing NanoPop-*bcd*3’UTR, MCP-GFP, and Vasa-GFP. GFP channel is shown. D. Translation of *suntag-nanos*-24xMS2 influenced by NanoPop_MCPGFP granules (top) and NanoPop_MCPGFP_VasaGFP (bottom) at the anterior. Zoom-in images are shown on the right. E. Quantification of the translation of *suntag-nanos*-24xMS2 at the anterior with various NanoPop granules. F. Quantification of the translation of *suntag-nanos-SREmut*-24xMS2 at the anterior with NanoPop_MCPGFP_VasaGFP granules. G. A proposed mechanism of translational activation by Vasa. Vasa helicase activity disrupts the structure of SRE in *nanos* 3’UTR, by which the binding of Smaug and translational repression is prevented.

To test whether RNA-unwinding activity was required for the translational activation by Vasa, we co-expressed NanoPop, MCPGFP, helicase mutant Vasa^E400Q^-GFP, and *suntag-nanos-MS2×24*. Localized mRNA in the designer granules showed no increase in translation, suggesting that helicase activity is indispensable for Vasa to activate translation of granule-localized mRNA (Fig. 3E and S3E). Next, we asked whether the translation activation is a unique function of Vasa, or a general capability of DEAD-box RNA helicase. We examined two DEAD-box RNA helicases, Belle (DDX3) and Me31B (DDX6), which both have function during germ cell development^52,53^. By expressing their GFP fusion proteins with NanoPop, MCPGFP, and *suntag-nanos-MS2×24,* we linked the designer NanoPop granules to Belle or ME31B. Neither helicase increased the translation of localized *suntag-nanos-MS2×24* mRNA *as* significantly as Vasa (Fig. 3E and Fig S3E). Next, we examined whether Vasa’s impact on *nanos* translation resulted from relieving Smaug’s repression. We generated Smaug resistant SRE-mutated RNA, *Suntag-nanos-SREmut-MS2×24*, and tested it in conjunction with the NanoPop_MCP-GFP_Vasa-GFP granule. *Suntag-nanos-SREmut-MS2×24* showed comparable level of translation to wildtype *suntag-nanos-MS2×24* localized in the designer granules, suggesting that the designer granule boosted translation mostly via counteracting repression by Smaug (Fig. 3F). Together these results suggest that Vasa, among DEAD box helicases, has the unique ability to activate the translation of RNP granule-localized mRNA through its RNA helicase activity. Potentially, Vasa unwinds the hairpin structure of the SRE and prevents Smaug from binding, thereby protecting *nanos* from repression (Fig. 3G).

### Validate the function of Vasa in native germ granules

After demonstrating Vasa’s role in activating translation in designer granules, we were curious to see whether Vasa is similarly involved in localized translation within native germ granules. To avoid the need for Vasa to assemble germ granules, we explored the possibility of acutely depleting Vasa in embryos after germ granule formation but before assessing translation activity. (Fig. 4A). We employed an auxin-inducible target protein degradation strategy, whereby embryos express Vasa tagged with an auxin-induced degron (AID) and GFP (Vasa-GFP-AID) along with TIR1 (Transport Inhibitor Response 1) protein, which targets the AID degron for polyubiquitination and proteasomal degradation^54,55^. After two hours of incubation with 0.5M auxin in a permeabilizing solution (see method), most embryos showed a strong decrease or complete ablation of the Vasa signal at the germplasm. However, *nanos* mRNA remained localized to the posterior pole, similar to untreated control embryos, suggesting that Vasa can be specifically depleted from germ granules post-granule formation without affecting mRNA localization (Fig. S4A). Although the protein degradation strategy was able to ablate Vasa from germplasm, it required a long incubation in a permeabilizing solution, which alone affected *nanos* translation activity in germplasm (Fig. S4B). Therefore, a more acute depletion of Vasa was needed.

**Figure 4.**
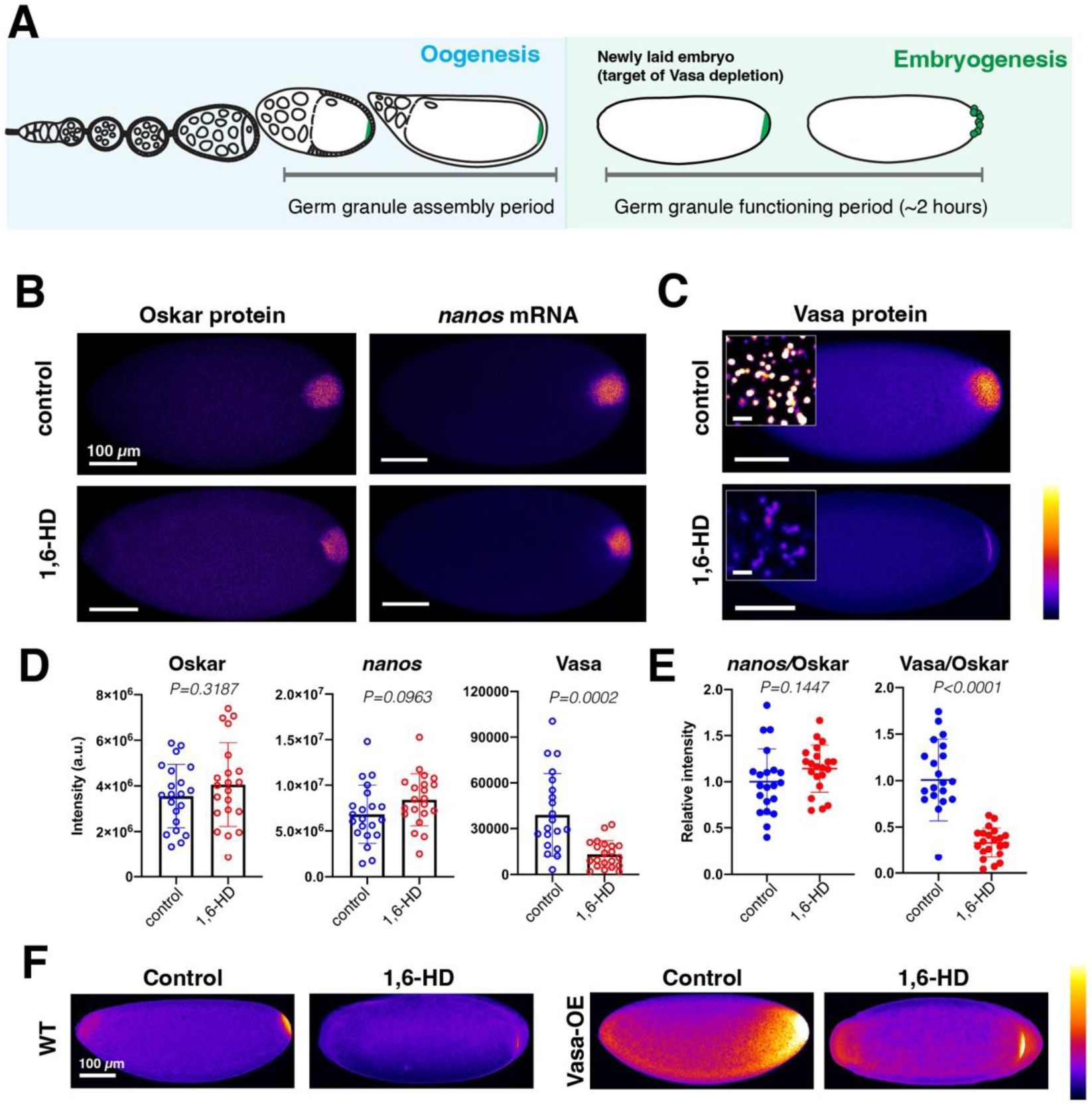
A. Schematics of Drosophila oogenesis when germ granule assembly occur, and early embryogenesis when germ granules function. Ideal time window to deplete Vasa is during early embryogenesis before pole cells form. B. Oskar protein (Oskar-GFP) and *nanos* mRNA (smFISH) were affected by treatment with 10% 1,6-HD for 20min. Fluorescence intensity is shown as a Fire LUT. C. Vasa protein (VasaGFP) was strongly depleted by treatment with 10% 1,6-HD for 20min. D. Quantification of fluorescence intensity of Oskar-GFP, Vasa-GFP, and *nanos* mRNA smFISH after control and 1,6-HD treatment. E. Quantification of *nanos* mRNA and Vasa level normalized by Oskar level. F. Vasa level (anti-Vasa staining) in WT embryos and *vasa-OE* embryos.

1,6-Hexanediol (1,6-HD) is an organic compound known to dissolve phase-separated biomolecular condensates by interfering with weak hydrophobic protein-protein or protein-RNA interactions^56^. We asked whether germ granules or select germ granule components could be disrupted by 1,6-HD. We incubated early Drosophila embryos with 10% 1,6-HD for 15 min, a condition expected to dissolve most liquid-like condensates, followed by fixation and staining. Oskar protein and *nanos* mRNA localization remained unchanged in germ granules after 1,6-HD treatment, suggesting that they have established stable interactions within germ granules (Fig. 4B). In contrast, Vasa protein was strongly depleted from germ granules after 1,6-HD treatment (Fig. 4C-E and Fig. S4C). Vasa has been shown to be recruited to germ granules through a protein-protein interaction between the Oskar LOTUS domain^27,28^, however other region in Oskar may also be also be able to recruit Vasa^34,57^, and Vasa can affect the biophysical properties of Oskar granules^58^. Given these close interactions of Vasa with Oskar, susceptibility to 1,6-HD was not necessarily expected. Significant depletion of Vasa by 1,6-HD implied that Vasa localization to germ granules involves weak, transient interactions, potentially via the intrinsically disordered region of Vasa. Vasa depletion phenotype could be rescued by over-expressing *vasa* (*vasa-OE*) and raising the overall concentration of Vasa inside embryos. The germplasm of *vasa-OE* embryos retained enriched Vasa signal even after 1,6-HD treatment (Fig. 4F). This supports the idea that Vasa localization in germ granules involves phase-separation-like condensation that is concentration-dependent^59^.

In addition to Oskar and *nanos* mRNA, other germ granule components, like Tudor protein and *germ cell-less* (*gcl*) mRNA, were also not affected by 1,6-HD treatment (Fig. 4SD and E). Germ granules’ morphology showed no obvious change by 1,6-HD either. Overall, Vasa is uniquely susceptible to depletion by 1,6-HD. After Vasa depletion, the remaining germ granule stayed largely intact. Thus, while Vasa is required for mRNA localization during germ granule formation, it is dispensable for maintaining mRNA within germ granules. This suggests that Vasa mediates RNA localization by passing the RNA, through its helicase activity, to the scaffold RNA-binding protein, Oskar. Localized RNA likely remained bound to Oskar and stabilized by RNA-RNA interaction, but not bound by Vasa.

Such specific and acute depletion of Vasa by 1,6-HD provided us with a valuable molecular handle to assess whether Vasa plays a role in activating translation in germ granules. We focused on *nanos* translation and used the well-characterized *suntag-nanos* to quantify translation in germplasm^40^. We found that following 1,6-HD treatment, significantly less *suntag-nanos* mRNA in germplasm was translated, and the intensities of translation spots, indicative of ribosome occupancy, significantly decreased (Fig. 5A-C). This result showed that 1,6-HD suppressed the translation of *nanos* in germplasm. Considering the potentially non-specific nature of the 1,6-HD effect, we asked whether 1,6-HD generally suppressed translation in the entire embryo. We used *suntag-nanos-tub3’UTR*, which has a *tubulin 3’UTR* instead of *nanos 3’UTR* and thus is translated throughout an embryo, reflecting general translation level^40^. Treatment of 1,6-HD did not lower the translation of *suntag-nanos-tub3’UTR*, suggesting that 1,6-HD specifically suppressed translation in germplasm, instead of inhibiting the overall translation level in embryos (Fig. S5A and B). Next, we investigated whether the decrease of *nanos* translation in germplasm was caused by Vasa depletion following 1,6-HD treatment. As Vasa depletion could be rescued by over-expressing *vasa* (*vasa-OE*), we asked whether Vasa-OE could rescue the translation reduction. Remarkably, translation of *suntag-nanos* was unperturbed in *vasa-OE* embryos following 1,6-HD treatment, suggesting that 1,6-HD suppressed *nanos* translation in germplasm specifically through depleting Vasa protein from germ granules (Fig. 5D-F). Moreover, 1,6-HD did not decrease the translation of *suntag-nanos-SREmut*, which is resistant to Smaug, both in germplasm and soma (Fig. 5G-I and Fig. S5C-D). This suggests that 1,6-HD specifically acted through Smaug to repress translation. Together, our data showed that Vasa activated the translation of *nanos* in germ granules by counteracting the repression posed by Smaug (Fig. 5J).

**Figure 5.**
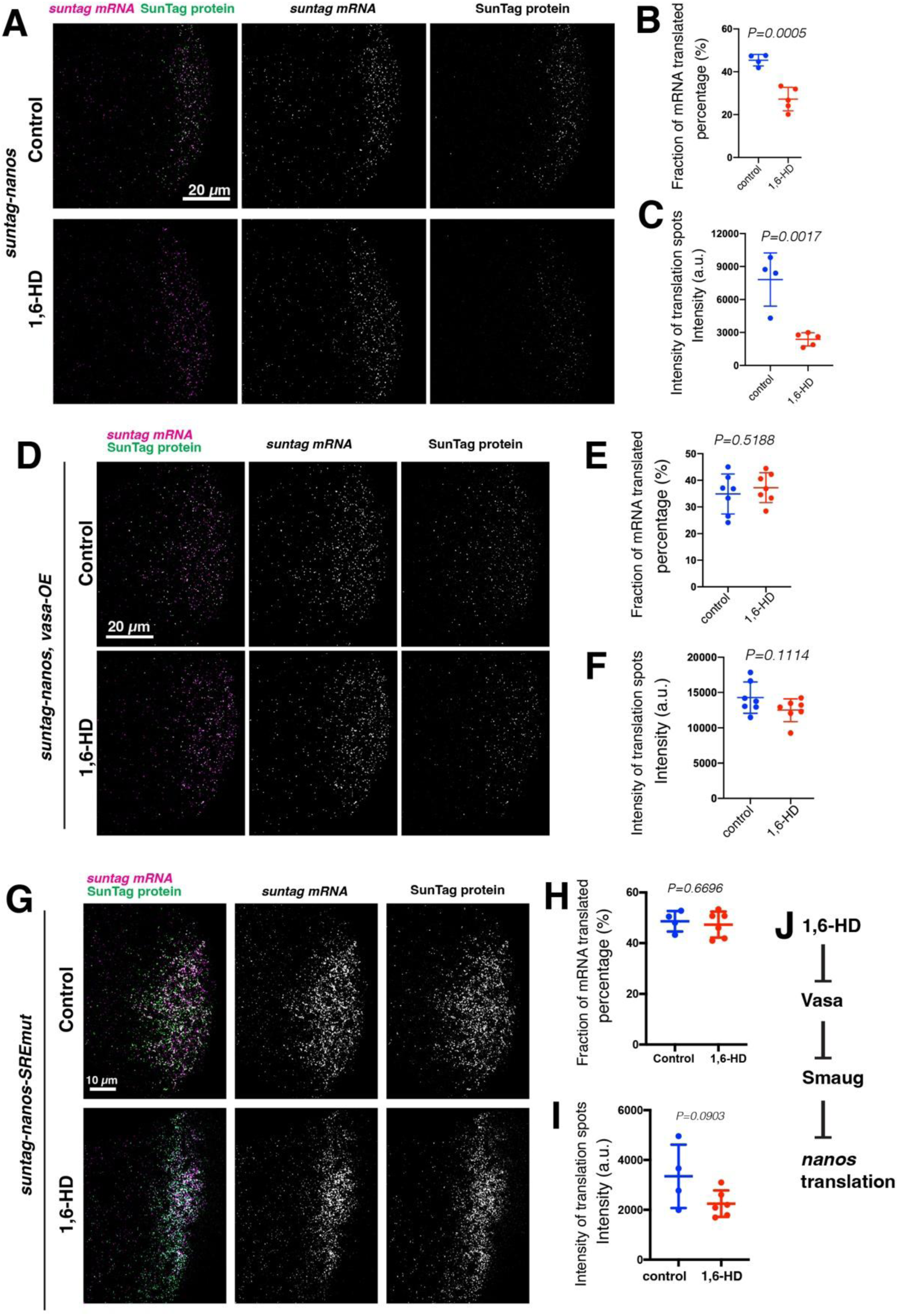
A-C. Translation of *suntag-nanos* in embryos treated with PBS (Control) and PBS with 10% 1,6-HD for 20min. Images of *suntag-nanos* mRNA (magenta) and SunTag protein spots (anti-GCN4, green) germplasm are shown in A. Translating fraction (B) and intensity of translation spots (C) are quantified. D-F. Translation of *suntag-nanos* in *vasa-OE* embryos treated with PBS (Control) and PBS with 10% 1,6-HD for 20min. Images are shown in D. Translating fraction (E) and intensity of translation spots (F) are quantified. G-I. Translation of *suntag-nanos-SREmut* in embryos treated with PBS (Control) and PBS with 10% 1,6-HD for 20min. Images are shown in G. Translating fraction (H) and intensity of translation spots (I) are quantified. J. Proposed mechanism of 1,6-HD effect and translation activation by Vasa. Vasa activates translation of *nanos* by blocking Smaug. 1,6-HD dissipates Vasa from germ granules so that Smaug can suppress *nanos* translation in germ granules.

## Discussion

As arguably the most conserved component of germ granules throughout evolution, the role of Vasa in germ granule assembly and function has been poorly established. Genetic analysis using Vasa mutant always complicated the interpretation due to the involvement of Vasa in various steps of germ granule biogenesis. Here, we established an *in vivo* reconstitution system in Drosophila germline and embryo utilizing self-assembly designer construct NanoPop. The system allowed us to test the role of individual proteins and/or domains in germ granule assembly and examine the effect of mutants without the complication of epistasis and pleiotropism. Using the system, we validated the role of RNA binding domain of Oskar in recruiting specific mRNA, and uncovered the necessity of Vasa and its RNA helicase activity in localizing mRNA. By circumventing the need of Vasa in RNA localization using MCP-MS2 system, we further demonstrated the necessity and sufficiency of Vasa in activating the translation of localized mRNA. An unexpected finding that Vasa can be acutely and specifically depleted by 1,6-HD from germ granules without affecting other essential components allowed us to validate our findings from the reconstituted system in native germ granules. With quantitative imaging of SunTag translation as a functional readout, we demonstrated that Vasa used its RNA unwinding activity to protect localized *nanos* from repression by Smaug.

Unwinding activity of RNA helicases has been posited to drive RNP granule disassembly by breaking up RNA-RNA interactions^4,5,60^. Our results suggest an opposite role for RNA helicases and their ability to unwind RNA: mediating RNA localization. The requirement of Vasa for *nanos* RNA localization was unexpected because Oskar has been considered the main RNA binder in germ granules and is supposedly able to recruit RNA by itself. Our results suggest that Oskar is necessary but not sufficient for RNA localization. It remains to be determined how Vasa and Oskar cooperate with one another to localize RNA in germ granules. Potentially, Vasa unwinds certain secondary structure in the RNA to expose the sequence element recognized by Oskar. The principle might also apply to other RNP granules: RNA unwinding by helicases precedes and is required for RNA interaction with the main RNA binders in the granule and RNA localization.

RNA helicases and their unwinding activity have been shown to promote translation. A classic case is that eIF4A and DDX3 unwind complex structures in the 5’UTR to allow translation initiation complex to scan through and find start codon^61–63^. YTHDC2 resolves secondary structures in coding sequences to facilitate translational elongation^64^. DDX3 can also promote translation within condensates by increasing molecular dynamics^60^. Our case of Vasa/DDX4 represents a new mechanism: an RNA helicase can protect RNA from its translational repressor. We speculate that Vasa does it by resolving the stem-loop structure of SRE in the *nanos* 3’UTR, which translational repressor Smaug recognizes and binds. Several conserved residues and motifs of Vasa have been implicated in translational activation with unclear mechanism^20,65^. Their corresponding mutants can be systematically tested using the NanoPop system to further dissect Vasa’s working mechanism.

Our study demonstrated a reductionist, bottom-up approach to studying RNP granules within their original cellular and physiological contexts. Until now, most understanding of biomolecular condensates has been derived from *in vitro* reconstitution using purified components, which is an elegant method for uncovering the principles behind condensate assembly and function. However, *in vitro* reconstitution faces significant challenges in replicating in vivo conditions and in purifying poorly-behaved disordered proteins. Our *in vivo* reconstitution, employing a synthetic biology approach, overcame these challenges by constructing and analyzing target condensates in their native environment, while avoiding epistatic effects that can complicate interpretation in traditional genetic studies. In addition to advancing our understanding of condensates, this synthetic biology strategy has opened new avenues for resolving various mechanistic questions in living organisms. Our work provides another example of the power of this innovative approach.

Limitation of the study: the synthetic granules made by NanoPop::OSK and VasaGFP only fulfilled part of germ granules’ functions. These granules could not induce pole cells and did not activate *nanos* translation as much as native germ granules. Further improvement is necessary to enhance RNA recruitment to localize full suite of mRNA required for pole cell formation. Vasa might not be the sole contributor of translational activation in germ granules. How other components are involved and their functional relationship with Vasa needs to be elucidated. In addition to *nanos*, numerous other mRNAs are also translated in germ granules. Whether and how Vasa influences their translation remains unclear. This work uncovered the role of Vasa helicase activity in antagonizing translational repressor Smaug but the exact mechanism remains to be clarified. Whether and how Vasa affects the structure of SRE and Smaug-*nanos* interaction requires the determination of *in vivo* structure and interactome of *nanos* mRNA and biochemical studies *in vitro*.

**Figure S1.**
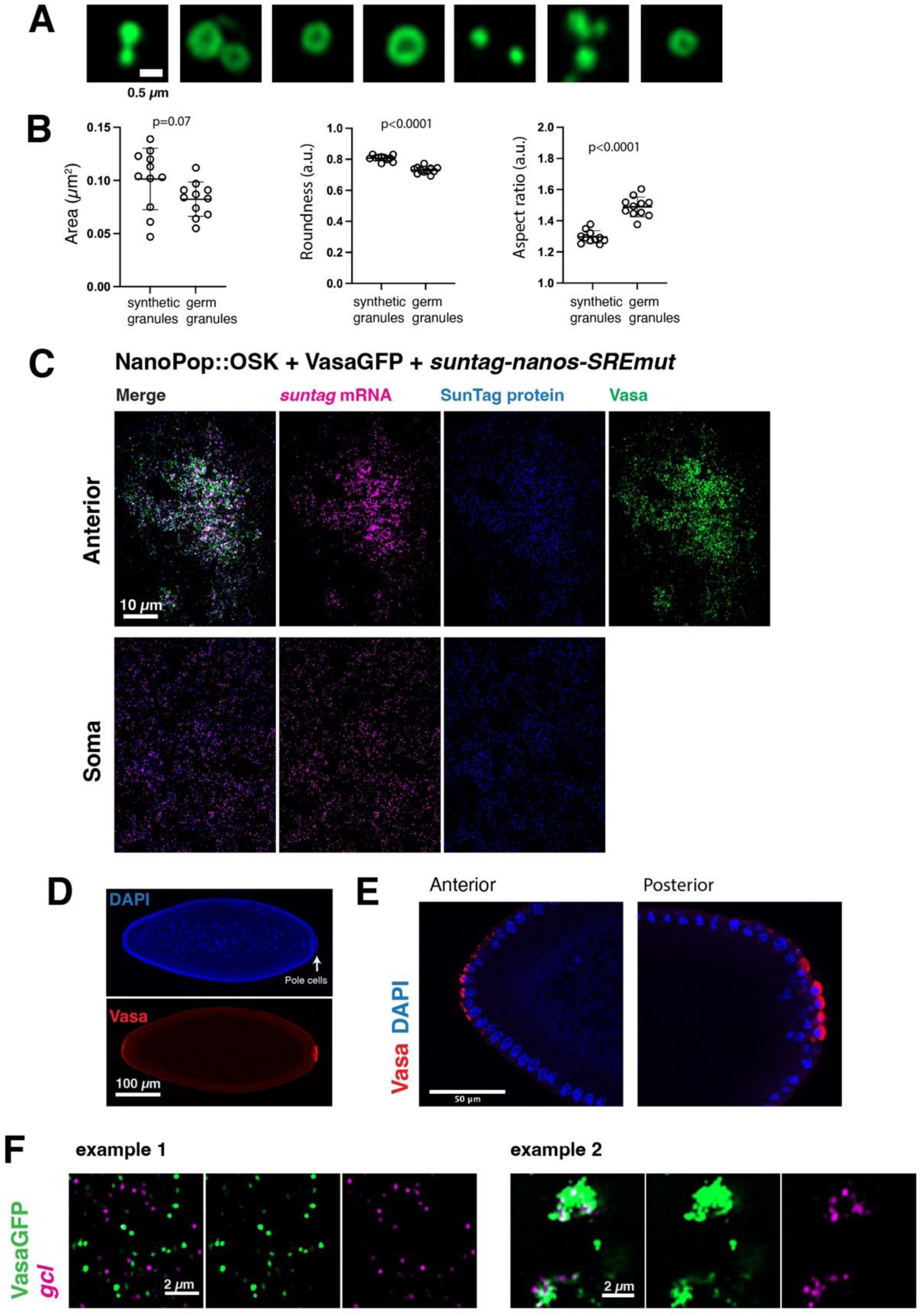
A. Gallery of synthetic NanoPop::OSK granules, labeled by VasaGFP, in a NanoPop::OSK_VasaGFP embryo. B. Comparing the size and morphology of synthetic NanoPop::OSK granules and native germ granules. Images of granules were segmented in Fiji and the area, roundness, aspect ratio (AR) of individual granules were measured. Each data point represents the average value of all quantified granules from one embryo. Roundness=1 or AR=1 means perfect circle. Higher AR means a more elongated shape. C. Translation of *suntag-nanos-SREmut* in the soma and the anterior of NanoPop::OSK_VasaGFP embryo, related to the quantification in Fig. 1I. D. A stage-5 NanoPop::OSK_VasaGFP embryo stained with anti-Vasa (red) and DAPI (blue). Pole cells (arrow) formed at the posterior pole but not anterior. E. Zoom-in views of the anterior and posterior poles of the embryo in B. Note the pole cells are filled with diffuse and strong Vasa staining (red). F. Rare localization of *gcl* mRNA (smFISH, magenta) to NanoPop::OSK granules (green). Example 1 represents the more prevalent situation where *gcl* mRNA did not localize. Example 2 shows a rare case when *gcl* localized to big, aggregated granules.

**Figure S2.**
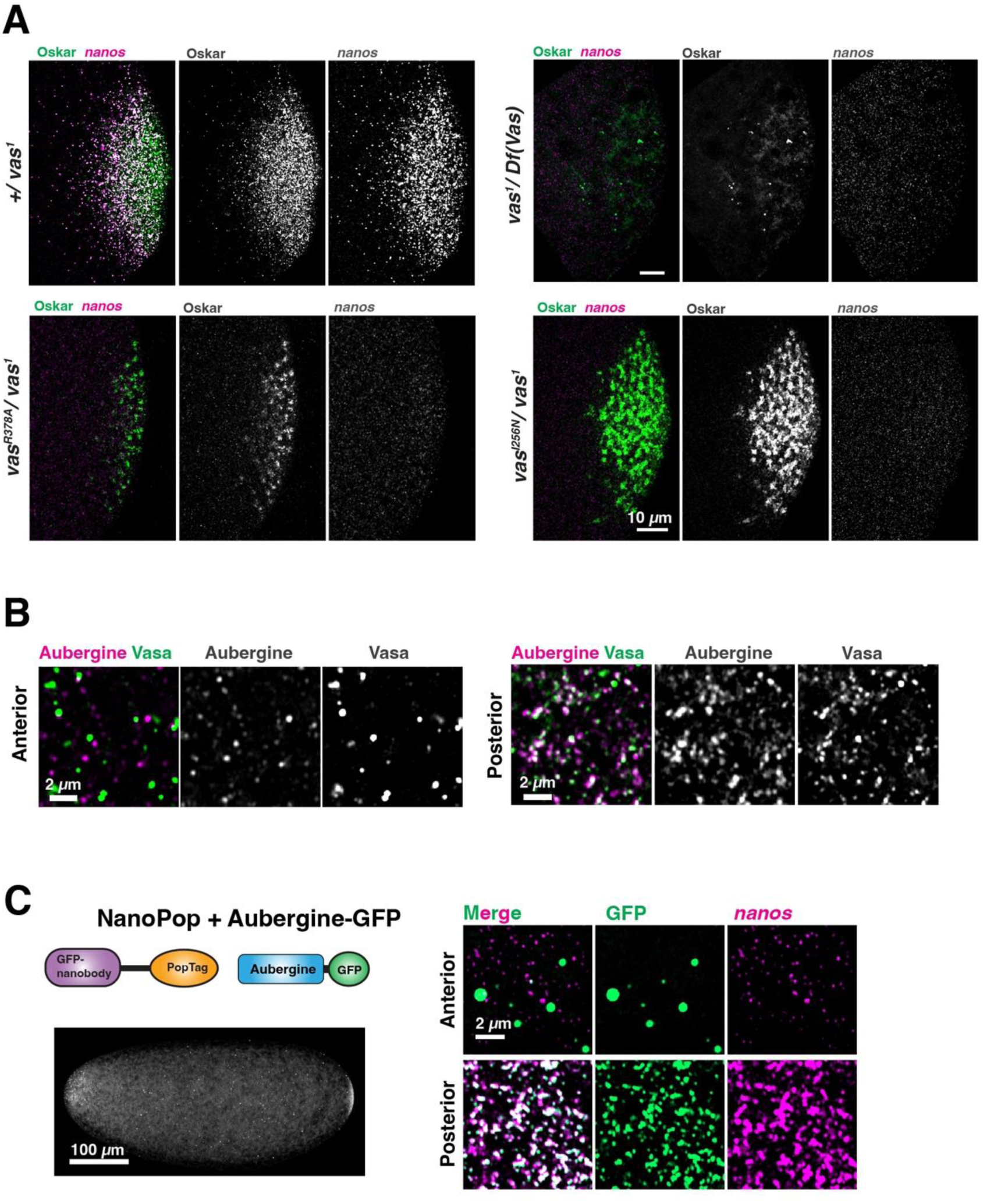
A. Localization of *nanos* (magenta) to native germ granules (Oskar, green) in WT (+/vas^1^), vasa null (vas^1^/Df(vas)), *vas^R^*^378^*^A^mApple* (*vas^R^*^378^*^A^mApple/vas*^1^), and *vas^I^*^256^*^N^* (*vas^I^*^256^*^N^/vas*^1^) embryos. B. The anterior and posterior poles of a NanoPop::OSK_VasaGFP embryo, stained with anti-Aub (magenta). Aub colocalized with Vasa (green) in the posterior germ granules but not in the anterior NanoPop::OSK granules. C. Embryos expressing NanoPop and GFP::Aub. *Nanos* mRNA (magenta) colocalized with Aub (green) at the posterior in germ granules. *Nanos* mRNA did not colocalize with Aub at the anterior in the NanoPop granules.

**Figure S3.**
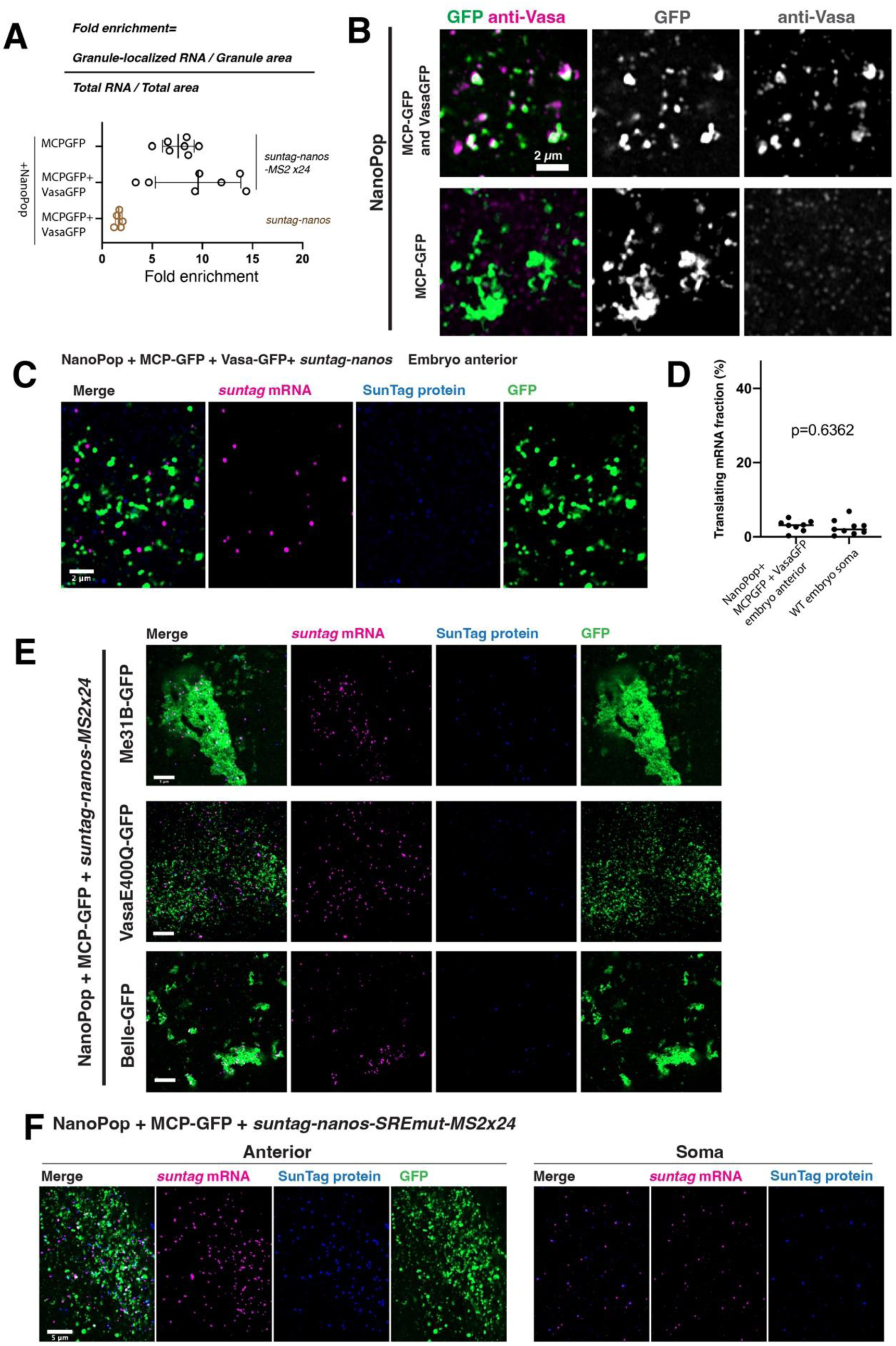
A. Quantification of mRNA localization: the enrichment of RNA density (RNA number divided by area) in synthetic granules over the entire embryo. NanoPop_MCPGFP and NanoPop_MCPGFP_VasaGFP granules showed 5 to 15-fold enrichment over the entire embryo (see Methods for quantification method). B. Vasa, detected by anti-Vasa (magenta), was localized to NanoPop_MCPGFP_VasaGFP granules but not NanoPop_MCPGFP granules (green). C. Image of *suntag-nanos* (no MS2) mRNA (magenta) and its translation (blue) with NanoPop_MCPGFP_VasaGFP granules. Note the lack of localization of mRNA and low translation. D. Quantification of translating fraction of *suntag-nanos* with NanoPop_MCPGFP_VasaGFP granules and *suntag-nanos* in the soma of WT embryos. E. Example images of *suntag-nanos-24xMS2* translation with NanoPop_MCPGFP_Me31BGFP, NanoPop_MCPGFP_Vasa^E400Q^GFP, and NanoPop_MCPGFP_BelleGFP granules. F. Example images of *suntag-nanos-SREmut-24xMS2* translation with NanoPop_MCPGFP_VasaGFP granules (anterior) and in soma.

**Figure S4.**
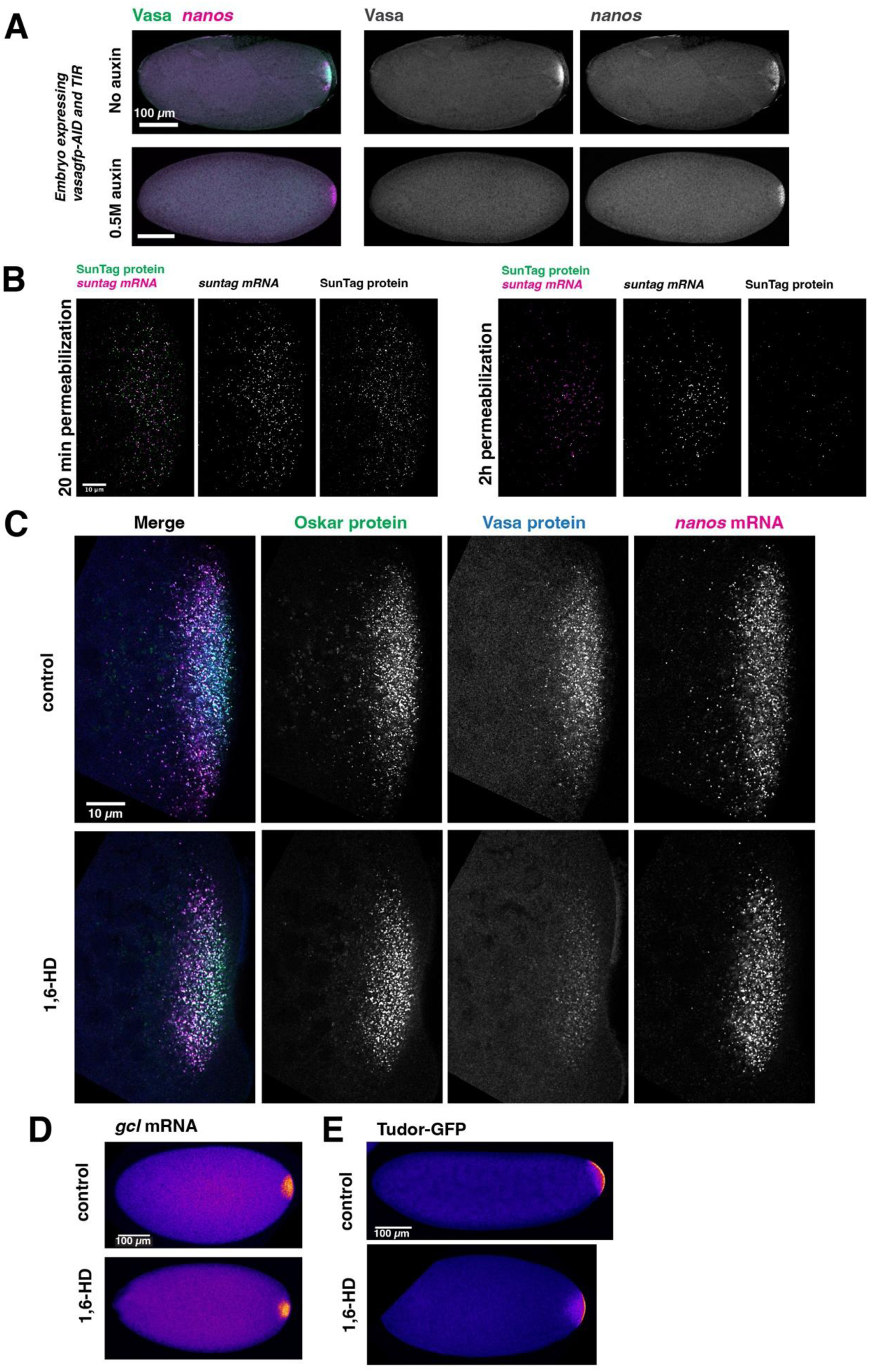
A. VasaGFP-AID embryos treated with 0.5M Auxin for 2h lost Vasa signal (green) but retained *nanos* mRNA (magenta) at the posterior. B. Two-hour permeabilization strongly reduced *suntag-nanos* translation in germplasm compared to 20-min permeabilization. C. Zoom-in views of germplasm of embryos treated with PBS (Control) and PBS with 10% 1,6-HD for 20min. Oskar, Vasa protein, and *nanos* mRNA are shown. D. Level of *gcl* mRNA (smFISH) in control and 1,6-HD treated embryos. E. Level of Tudor-GFP in control and 1,6-HD treated embryos.

**Figure S5.**
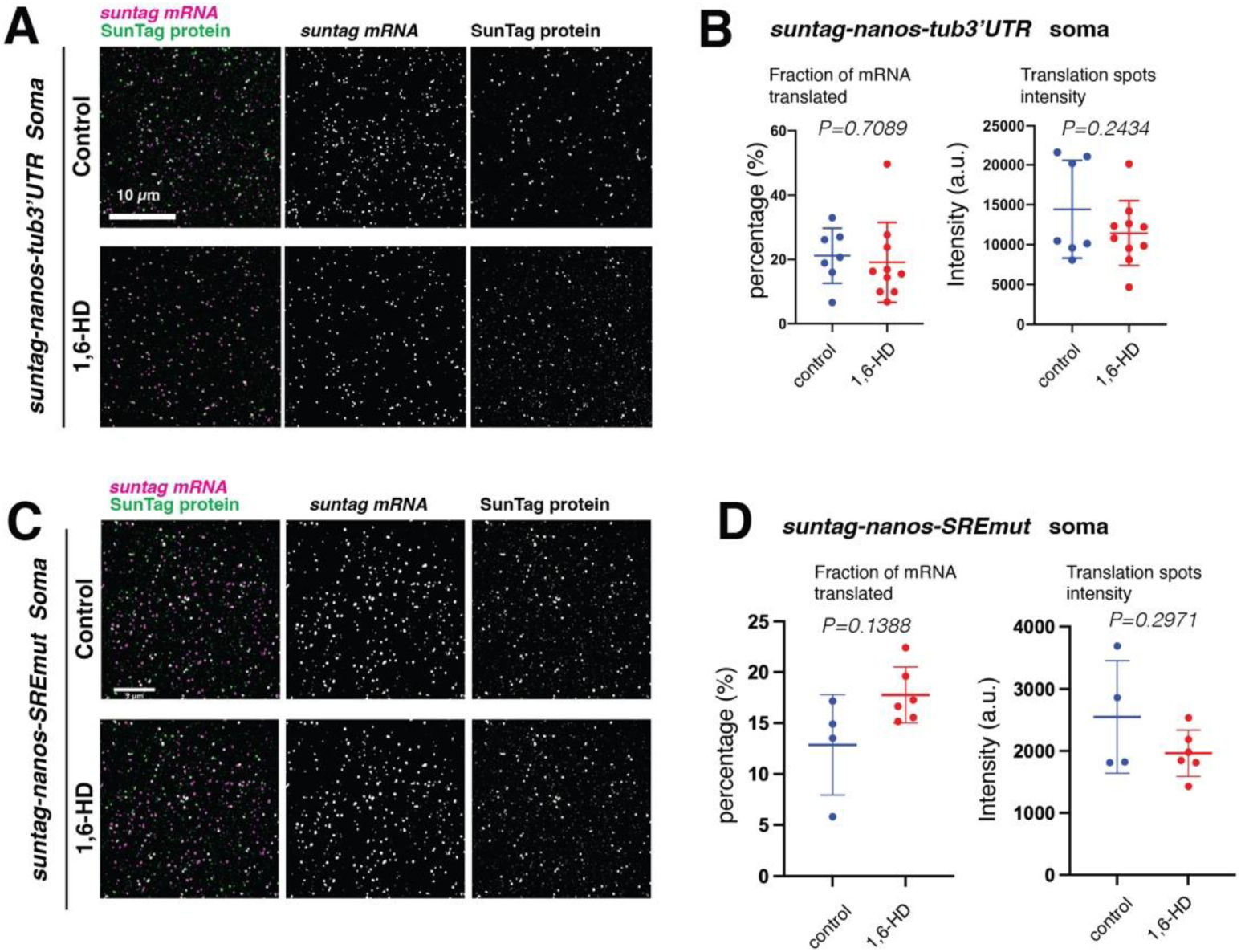
A-B. Translation of *suntag-nanos-tub3’UTR* in embryo soma treated with PBS (Control) and PBS with 10% 1,6-HD for 20min. C-D. Translation of *suntag-nanos-SREmut* in embryo soma treated with PBS (Control) and PBS with 10% 1,6-HD for 20min.

## Materials and methods

### Fly stocks and genetics

Fly stocks were maintained at 25°C. Detailed experimental genotypes and sources of fly stocks are listed in Supplementary Table 1.

### Cloning, transgenesis, and CRISPR

All the constructs were made using In-Fusion cloning (Takara Bio) unless specified otherwise. All PCR was performed using CloneAmp HiFi PCR premix (Takara Bio #639298). The sequences and maps of all NanoPop constructs were provided in the supplemental file. The plasmids and full sequence files will be provided upon request. For transgenesis, individual plasmids were injected into attP2 or attP40 lines (the BestGene Inc.) and transformants were screened for the presence of the *mini-white* eye markers.

#### UAS-NanoPop-bcd3’UTR

NanoPop sequence was a kind gift from Keren Lasker. This construct contains an anti-GFP nanobody at the N-terminus and a Poptag at the C-terminus, with a 102-amino acid linker in the middle. This construct was PCR-amplified, inserted into a UASz vector^66^, and appended with a *bcd*3’UTR.

#### UAS-NanoPop::OSK-bcd3’UTR

The OSK domain sequence of Oskar protein (TPTP…TSLEY) was PCR-amplified and inserted after a (GGGS)x3 linker that follows anti-GFP nanobody. OSK sequence is followed by the rest of the linker sequence (87 amino acids) and the Poptag.

#### UAS-NanoPop::G3BP1-bcd3’UTR

The C-terminal RRM/RGG rich region of human G3BP1 protein (RHPD…APRQ) was PCR-amplified from a G3BP1 plasmid (addgene #119950), inserted after anti-GFP nanobody and a (GGGS)x3 linker, and followed by a 7-amino acids linker and the Poptag.

#### VasaE400Q-GFP

A sequence containing *vasa* promoter, coding sequence, and 3’UTR was PCR-amplified from the genomic DNA of a GFP-Vasa transgenic strain^53^ and a vector derived from UASz, in which the UAS sequence, 5’ and 3’UTR were removed. The E400Q mutation was introduced by PCR with a primer pair containing the mutation.

#### UAS-suntag-nanos-24xMS2 and UAS-suntag-nanos-SREmut-24xMS2

24xMS2v5 sequence was PCR-amplified from a MS2-containing plasmid (addgene #119946) and inserted into a *UAS-suntag-nanos* plasmid^40^. The 24xMS2v5 was placed after the stop codon of nanos coding sequence and followed by nanos 3’UTR. To introduce mutations at the two SRE sequences in the *nanos* 3’UTR, a gBlock of the sequence containing the mutant SREs was synthesized and replaced the WT sequence in the *UAS-suntag-nanos* plasmid. The wild-type sequences are SRE1: GCAGAGGCT<u>C</u>T<u>G</u>GCAGCTTTTGC, and SRE2: AAATAGCGC<u>C</u>T<u>G</u>GCGCGTTCGAT. The mutant has underlined C mutated to G, and underlined G mutated to C.

*VasaR378A-mApple* was generated by homology-directed recombination following CRISPR/Cas9 gene targeting. Endogenous *vasa* locus was targeted using two guide RNAs: (guide#1: CGCCACTCCGGGACGACTTC; guide#2: AGCAATGGGATTGAAATGTA), which were cloned into pCFD-dU6:3gRNA (DGRC_1362). For recombination template, part of *vasa-mApple* sequence was PCR-amplified from genomic DNA of *vasa-mApple* CRISPR line (DGRC#118617) and inserted into pScarless-HD-DsRed (Addgene #64703). The R378A mutation was introduced using PCR with a primer pair that contains the mutation. Plasmids of the recombination template and two guide RNAs were injected into a Cas9-expressing line (BDSC#51324). Transformants were screened using the DsRed eye marker. DsRed was removed from the *VasaR378A-mApple* allele by crossing with a line expressing PiggyBac transposase in germline.

### Design and expression of NanoPop constructs

NanoPop constructs were expressed at the anterior of the embryo using *bcd3’UTR*, which targets the mRNA to the anterior pole of oocytes and resultant embryos. The same strategy has been used to express Oskar anteriorly and induce functional germ granules^18^. The constructs were under the control of a UAS element and were driven by a maternally-expressed α-tubulin GAL4VP16 (MatαGAL4) which expresses in ovaries during mid-oogenesis, a similar stage when native germ granules biogenesis occurs. VasaGFP, GFP, and other GFP-fusion proteins were also maternally expressed in ovaries and deposited into embryos to allow synthetic NanoPop granule assembly. Synthetic granules were visualized through GFP or GFP-fusion proteins.

Two different strains expressing GFP-fused Vasa were used. One is the GFP CRISPR knock-in at the C-terminus of the endogenous Vasa^67^, the other one is a GFP-Vasa (N-terminally tagged) transgene on the 3rd chromosome^53^. Both GFP-fused Vasa alleles were able to functionally replace wild-type Vasa. And two VasaGFP constructs formed synthetic granules with NanoPop constructs with similar efficiency, morphology, and functionality. Which allele being used was decided based on the chromosome availability in crossings.

Embryos expressing *NanoPop::OSK-bcd3’UTR* or *NanoPop-bcd3’UTR* alone developed normally. However, we found that co-expressing VasaGFP with *NanoPop::OSK-bcd3’UTR* or *NanoPop-bcd3’UTR* caused lethality in embryos. Very few embryos finished gastrulation. This effect appeared to be specific to VasaGFP but not other GFP-fusion proteins co-expressed with NanoPop constructs. The cause of this synthetic lethality is being investigated.

### Embryo collection and fixation

Drosophila embryos were collected for 3h on an apple juice plate, dechorionated by incubating with 50% bleach solution for two minutes, extensively washed, and transferred to a scintillation vial containing a 1:1 (v/v) mixture of heptane and 4% paraformaldehyde in PBS (phosphate-buffered saline), in which embryos were permeabilized and fixed for 20 min. The paraformaldehyde was removed with a Pasteur pipette, followed by adding methanol and vigorous shaking for 15 s to remove the vitelline membrane. Embryos were washed three times for 5 min with methanol before being stored at 4°C in methanol.

### Immunofluorescence

Embryos were rehydrated by washing for 5 min with 50% methanol with PBS-Triton 0.3% and then washed and permeabilized for 3x 15 min in PBS-TritonX-100 0.3%. Embryos were blocked with 1% BSA in PBS-TritonX-100 0.3% for 30 min and subsequently incubated with primary antibodies diluted in the blocking solution overnight at 4°C.Embryos were washed five times for 10min with PBS-TritonX-100 0.3%, blocked for 30min, and incubated with secondary antibodies with 1:1000 dilution for 4 h at room temperature. Then embryos were washed five times for 10 min with PBS-Triton 0.3%, stained with DAPI, and mounted with ProLong Glass mounting medium (ThermoFisher, P36980).

### Single-molecule RNA fluorescent in situ hybridization (smFISH)

Stellaris RNA FISH probes against *suntag*, *nanos*, *pgc*, and *gcl* sequences were used for hybridization. To perform smFISH on fixed embryos, stored embryos were rehydrated by washing for 5 minutes with 50% methanol with PBS-Tween 0.1% and washing three times for 5 min in PBS-Tween 0.1%. Embryos were then washed with a pre-hybridization buffer containing 2xSSC and 10% formamide (Fisher Scientific, AM9342) for 10 minutes at room temperature. The embryos were then incubated at 37°C for 3h in the hybridization mix (60µL hybridization mix per sample with 50-100 embryos) containing 2xSSC, 10% (v/v) deionized formamide, 0.1 mg/ml E.coli tRNA, 0.1mg/ml salmon sperm DNA, 10mM Ribonucleoside Vanadyl Complex (NEB, S1402S), 2mg/ml BSA, 80ng Stellaris probes and 10% (v/v) Dextran sulfate. After hybridization, embryos were washed with the pre-hybridization buffer twice for 15 minutes at 37°C. The embryos were washed with PBS-Tween 0.1% three times for 5min, stained with DAPI, and mounted with ProLong Glass mounting medium.

When anti-GCN4 is used to detect SunTag protein, IF was performed after smFISH. Following the 2×15min washes with pre-hybridization buffer, embryos were washed and permeabilized with PBS-TritonX-100 0.3% for 45 minutes. Embryos were blocked with 1% BSA in PBS-TritonX-100 0.3% for 30 minutes and then incubated with rabbit anti-GCN4 (Novus Bio.) with 1:1000 dilution overnight at 4°C. Embryos were washed with PBS-TritonX-100 0.3% five times for 10 minutes, blocked for 30min, and incubated with anti-rabbit secondary antibody (1:1000) for 4h at room temperature. Embryos were then washed with PBS-Triton 0.3% five times for 10 min, stained with DAPI, and mounted with ProLong Glass mounting medium.

### Embryo permeabilization and incubation with compounds

Drosophila embryos were dechorionated, washed, and transferred to a scintillation vial containing a 1:1 (v/v) mixture of heptane and PBS (control). For AID-mediated protein degradation, auxin was added to PBS. For acute Vasa depletion, 1,6-hexanediol (1,6-HD) was added to PBS. After incubation (∼2 hours for auxin, 20 min for 1,6-HD), PFA was added to the vial to reach a final concentration of 4% in the aqueous phase. After 20min fixation, the aqueous phase was removed, followed by adding methanol and vigorous shaking for 15 s to remove the vitelline membrane. Embryos were washed three times for 5 min with methanol before being stored at 4°C in methanol.

### Microscopy

Microscopic images were acquired using Zeiss LSM980 confocal microscope. Images of whole embryos were acquired using 10x 0.3 Numerical Aperture (NA) air objective. High-resolution images were acquired using Plan-Apochromat 63x /1.4NA oil objective with AiryScan 2 detector and SR mode. DAPI was excited by a 405nm laser. Red fluorophores (mApple, mCherry, Alexa Fluor 555, or Alexa Fluor 561) were excited by a 561nm laser. Green fluorophores (GFP, YFP, or Alexa Fluor 488) were excited by a 488nm laser. The far-red fluorophore (Alexa Fluor 647) was excited by a 639nm laser. Imaging acquisition and processing settings have been described in a previous study^40^.

### Translation quantification

Method of quantifying translation activity using SunTag foci has been described in detail in a previous study^40^. In brief, the quantification was performed using MATLAB-based software FISH-Quant_v3, in which the foci of *suntag* mRNA (smFISH) and SunTag protein (anti-GCN4) were detected and 3D coordinates and intensities of individual foci were determined. The co-localization between the detected mRNA and protein foci was analyzed by the DualColor program of FISH-Quant with 400 nm as the maximum distance between two spots to be considered co-localized. This analysis provides the percentage of mRNA foci co-localized with protein foci, which represents the percentage of translating mRNA. The intensity of SunTag protein correlates with ribosome occupancy.

### Image quantification

The abundance of specific protein or mRNA (Oskar, Vasa, Tudor, *nanos*, and *gcl*) in germplasm was quantified using whole-embryo images in Fiji. The germplasm area was selected based on the enriched signal of the germplasm marker Oskar, which remained unchanged after 1,6-HD treatment. The integrated signal of the area of the target channel was measured. Next, the same region of interest (ROI) was moved to a nearby somatic region and the integrated intensity was measured as background, which was subtracted from the germplasm signal.

To quantify the size and morphology of germ granules and synthetic granules, a single z-slice 63x high resolution image of granules (GFP channel) was used. A threshold was applied to the image using the default threshold setting in Fiji. Next, the area, roundness, and aspect ratio of individual granules were measured using the Analyse Particles function. The average values were taken to represent the granules of individual embryos.

### RNA localization

RNA localization to germ granules and synthetic granules were quantified using two different methods. For *nanos* localization, Pearson Correlation Coefficient (PCC) between *nanos* signal and GFP (granule) signal was measured using the BIOP-JACoP plug-in of Fiji program. Thresholding was automatic and the thresholding methods do not affect PCC outcomes. For suntag-nanos-24xMS2, which was lowly expressed and individual mRNA can be resolved inside granules, its localization to NanoPop_MCPGFP granules was measured as the enrichment level inside the granules. Specifically, images of embryo anterior containing NanoPop granules taken with 63x objective were used. The mRNA spots were identified and counted using the ComDet v0.5.5 plug-in in Fiji from the entire field of view and from granule areas, which were defined using thresholding tools in Fiji. The overall density of mRNA in the embryo was determined by normalizing the total number of identified mRNA with the area of the embryo in the image. The density of mRNA in granules was determined by normalizing the number of granule-localized mRNA with the area of granules. Fold enrichment is calculated as granule mRNA density divided by overall density.

## Supporting information

Supplemental sequence file

## Acknowledgement

We thank past and present members of the Lehmann lab for helpful suggestions throughout this work. We thank Sanaya Iyer for assistance with data acquisition. We thank Keren Lasker for kindly sending us NanoPop plasmids. We thank the Bloomington Drosophila Stock Center (NIH P40OD018537), Kyoto Drosophila Stock Center, and the Vienna Genome Resource Center, as well as Akira Nakamura for fly stocks and the Drosophila Genome Resource Center (NIH 2P40OD010949) for reagents. We are grateful to the FlyBase consortium for providing data and curators (NHGRI/NIH U41HG000739 and U24HG013300). This work was, in part, supported by NIH award (5R01HD110546). R.C. is supported by a Jerome and Florence Brill Graduate Student Fellowship. J.S.G. is supported by NIH F32 award (1F32HD111257). H.Z. is supported by NSF GRFP under Grant No. 2141064 and Dean of Science fellowship (MIT).

## Author contributions

R.C. and R.L. designed the experiments. R.C., H.Z., and J.S.G. performed the investigation. R.C. wrote the original draft. All authors reviewed and edited the text. R.L. acquired the funding. R.L. supervised the work.

## Data and materials availability

All data needed to evaluate the conclusions in the paper are present in the paper and/or the Supplementary Materials.

