## Supplemental sequence file for "Synthetic germ granules reveal a direct role of Vasa/DDX4 in RNA localization and translational activation"

Construct sequences:

UAS-NanoPop-bcd3’UTR. (Nanobody PopTag)

CGGAGTACTGTCCTCCGAGCGGAGTACTGTCCTCCGAGCGGAGTACTGTCCTCCGAGCGGAGTACTGTCCTCCGAGCGGAGTACTGTCCTCCGAGCGGAGACTCTAGCCCTAGGGCATGCCTGCAGGTCGGAGTACTGTCCTCCGAGCGGAGTACTGTCCTCCGAGCGGAGTACTGTCCTCCGAGCGGAGTACTGTCCTCCGAGCGGAGTACTGTCCTCCGAGCGGAGACTCTAGCGCTAGCGACGTCGAGCGCCGGAGTATAAATAGAGGCGCTTCGTCTACGGAGCGACAATTCAATTCAAACAAGCAAAGATCTAAAAGGTAGGTTCAACCACTGATGCCTAGGCACACCGAAACGACTAACCCTAATTCTTATCCTTTACTTCAGGGGCCGCAACTTAAAAAAAAAAATCAAAGGATCCCATGgttcaactggtggaaagcggcggtgctctggtacaaccgggcggtagtctgcgcctgagctgtgccgcaagcggtttcccagtcaaccgctactctatgcgttggtatcgccaggcgcctggtaaagaacgtgaatgggttgccggcatgagcagtgcgggcgatcgttctagttacgaggactctgttaaaggtcgttttacaattagccgtgatgatgcgcgcaataccgtgtatctgcaaatgaacagtctgaagccggaggacaccgcagtatattattgcaatgtcaacgtggggtttgaatattggggccaggggactcaggtgacggtgagctctggaggcggtggttccggaggcggcggctccggaggaggggggtcaccagctggcgccctgatcaacatgcacggcacagatgcccccgctgagccggcagctgaagcagcgcccccaccgccaccagaaccggaacccgagcctgtgagctttgatgacgaggtgctcgaattgacggaccctatagcgcctgagcctgagctgccacccctggagactggcgatattgacgtgtattccccacccgagcctgaatccgagcccgcctacaccccaccccctgcagcccccgtatttgatcgcgatgaggttgctgaacagctggtgggggtgtccgctgcttctgctgcggcctctgccttcgggtcattatccagcgcacttttgatgccaaaagatgggcgcactctggaggatgtggtgagagaactgctgcgtcctctgcttaaagaatggcttgaccaaaacctcccgaggatcgtcgagacaaaagtagaggaggaggtgcagcgcatctcccgcggacgcggcgcaTAACCTGGATGAGAGGCGTGTTAGAGAGTTTCATTAGCTTTAGGTTGGACTAGTCCATCTACGCGTAGAAAGTTAGGTCTAGTCCTAAGATCCGTGTAAATGGTTCCCAGGGAAGTTTTATGTACTAGCCTAGTCAGCAGGCCGCACGGATTCCAGTGCATATCTTAGTGATACTCCAGTTAACTCTATACTTTCCCTGCAATACGCTATTCGCCTTAGATGTATCTGGGTGGCTGCTCCACTAAAGCCCGGGAATATGCAACCAGTTACATTTGAGGCCATTTGGGCTTAAGCGTATTCCATGGAAAGTTATCGTCCCACATTTCGGAAATTATATTCCGAGCCAGCAAGAAAATCTTCTCTGTTACAATTTGACATAGCTAAAAACTGTACTAATCAAAATGAAAAATGTTTCTCTTGGGCGTAATCTCATACAATGATTACCCTTAAAGATCGAACATTTAAACAATAATATTTGATATGATATTTTCAATTTCTATGCTATGCCAAAGTGTCTGACATAATCAAACATTTGCGCATTCTTTGACCAAGAATAGTCAGCAAATTGTATTTTCAATCAATGCAGACCATTTGTTTCAGATTCGGAGATTTTTTGCTGCCAAACGGAATAACTATCATAGCTCACATTCTATTTACATCACTAAGAAGAGCATTGCAATCTGTTAGGCCTCAAGTTTAATTTTAAAATGCTGCACCTTTGATGTTGTCTCTTTAAGCTTTGTATTTTTAATTACGAAAATATATAAGAACTACTCTACTCGGGTAAATTGTGACTAACTAC

UAS-NanoPop::OSK-bcd3’UTR. (Nanobody OSK PopTag)

CGGAGTACTGTCCTCCGAGCGGAGTACTGTCCTCCGAGCGGAGTACTGTCCTCCGAGCGGAGTACTGTCCTCCGAGCGGAGTACTGTCCTCCGAGCGGAGACTCTAGCCCTAGGGCATGCCTGCAGGTCGGAGTACTGTCCTCCGAGCGGAGTACTGTCCTCCGAGCGGAGTACTGTCCTCCGAGCGGAGTACTGTCCTCCGAGCGGAGTACTGTCCTCCGAGCGGAGACTCTAGCGCTAGCGACGTCGAGCGCCGGAGTATAAATAGAGGCGCTTCGTCTACGGAGCGACAATTCAATTCAAACAAGCAAAGATCTAAAAGGTAGGTTCAACCACTGATGCCTAGGCACACCGAAACGACTAACCCTAATTCTTATCCTTTACTTCAGGGGCCGCAACTTAAAAAAAAAAATCAAAGGATCCCATGgttcaactggtggaaagcggcggtgctctggtacaaccgggcggtagtctgcgcctgagctgtgccgcaagcggtttcccagtcaaccgctactctatgcgttggtatcgccaggcgcctggtaaagaacgtgaatgggttgccggcatgagcagtgcgggcgatcgttctagttacgaggactctgttaaaggtcgttttacaattagccgtgatgatgcgcgcaataccgtgtatctgcaaatgaacagtctgaagccggaggacaccgcagtatattattgcaatgtcaacgtggggtttgaatattggggccaggggactcaggtgacggtgagctctggaggcggtggttccggaggcggcggctccggaggaggggggtcaACGCCCACGCCAACGATTTTAACCAGTGGAACCTACAACGATTCTCTGCTGACGATTAACTCGGATTACGATGCCTATCTGCTGGACTTTCCGCTTATGGGCGATGATTTTATGCTATATCTCGCCCGAATGGAGCTAAAATGCCGATTTAGGCGTCACGAACGCGTCCTGCAGTCAGGACTTTGTGTATCCGGACTGACGATCAATGGCGCCCGAAATCGTTTAAAAAGAGTCCAATTACCCGAGGGTACTCAGATCATCGTCAATATCGGATCGGTGGACATTATGCGCGGCAAGCCTTTGGTTCAGATCGAGCACGATTTTCGGCTACTGATCAAGGAGATGCACAATATGCGATTGGTGCCGATTCTAACAAATCTTGCACCGCTGGGCAACTATTGTCACGATAAGGTATTATGTGACAAAATCTACCGATTCAACAAGTTTATCCGAAGCGAATGCTGTCACCTAAAGGTCATCGACATACACTCCTGTCTAATCAACGAAAGGGGCGTGGTGCGATTTGATTGCTTTCAGGCCTCACCACGCCAAGTCACCGGTTCCAAGGAACCCTATCTGTTCTGGAACAAAATCGGTCGGCAGCGCGTACTGCAAGTTATTGAAACGAGTCTGGAGTATccagctggcgccctgatcaacatgcacggcacagatgcccccgctgagccggcagctgaagcagcgcccccaccgccaccagaaccggaacccgagcctgtgagctttgatgacgaggtgctcgaattgacggaccctatagcgcctgagcctgagctgccacccctggagactggcgatattgacgtgtattccccacccgagcctgaatccgagcccgcctacaccccaccccctgcagcccccgtatttgatcgcgatgaggttgctgaacagctggtgggggtgtccgctgcttctgctgcggcctctgccttcgggtcattatccagcgcacttttgatgccaaaagatgggcgcactctggaggatgtggtgagagaactgctgcgtcctctgcttaaagaatggcttgaccaaaacctcccgaggatcgtcgagacaaaagtagaggaggaggtgcagcgcatctcccgcggacgcggcgcaTAACCTGGATGAGAGGCGTGTTAGAGAGTTTCATTAGCTTTAGGTTGGACTAGTCCATCTACGCGTAGAAAGTTAGGTCTAGTCCTAAGATCCGTGTAAATGGTTCCCAGGGAAGTTTTATGTACTAGCCTAGTCAGCAGGCCGCACGGATTCCAGTGCATATCTTAGTGATACTCCAGTTAACTCTATACTTTCCCTGCAATACGCTATTCGCCTTAGATGTATCTGGGTGGCTGCTCCACTAAAGCCCGGGAATATGCAACCAGTTACATTTGAGGCCATTTGGGCTTAAGCGTATTCCATGGAAAGTTATCGTCCCACATTTCGGAAATTATATTCCGAGCCAGCAAGAAAATCTTCTCTGTTACAATTTGACATAGCTAAAAACTGTACTAATCAAAATGAAAAATGTTTCTCTTGGGCGTAATCTCATACAATGATTACCCTTAAAGATCGAACATTTAAACAATAATATTTGATATGATATTTTCAATTTCTATGCTATGCCAAAGTGTCTGACATAATCAAACATTTGCGCATTCTTTGACCAAGAATAGTCAGCAAATTGTATTTTCAATCAATGCAGACCATTTGTTTCAGATTCGGAGATTTTTTGCTGCCAAACGGAATAACTATCATAGCTCACATTCTATTTACATCACTAAGAAGAGCATTGCAATCTGTTAGGCCTCAAGTTTAATTTTAAAATGCTGCACCTTTGATGTTGTCTCTTTAAGCTTTGTATTTTTAATTACGAAAATATATAAGAACTACTCTACTCGGGTAAATTGTGACTAACTAC

UAS-NanoPop::OSK-bcd3’UTR. (Nanobody G3BP1RBD PopTag)

CGGAGTACTGTCCTCCGAGCGGAGTACTGTCCTCCGAGCGGAGTACTGTCCTCCGAGCGGAGTACTGTCCTCCGAGCGGAGTACTGTCCTCCGAGCGGAGACTCTAGCCCTAGGGCATGCCTGCAGGTCGGAGTACTGTCCTCCGAGCGGAGTACTGTCCTCCGAGCGGAGTACTGTCCTCCGAGCGGAGTACTGTCCTCCGAGCGGAGTACTGTCCTCCGAGCGGAGACTCTAGCGCTAGCGACGTCGAGCGCCGGAGTATAAATAGAGGCGCTTCGTCTACGGAGCGACAATTCAATTCAAACAAGCAAAGATCTAAAAGGTAGGTTCAACCACTGATGCCTAGGCACACCGAAACGACTAACCCTAATTCTTATCCTTTACTTCAGGGGCCGCAACTTAAAAAAAAAAATCAAAGGATCCCATGgttcaactggtggaaagcggcggtgctctggtacaaccgggcggtagtctgcgcctgagctgtgccgcaagcggtttcccagtcaaccgctactctatgcgttggtatcgccaggcgcctggtaaagaacgtgaatgggttgccggcatgagcagtgcgggcgatcgttctagttacgaggactctgttaaaggtcgttttacaattagccgtgatgatgcgcgcaataccgtgtatctgcaaatgaacagtctgaagccggaggacaccgcagtatattattgcaatgtcaacgtggggtttgaatattggggccaggggactcaggtgacggtgagctctggaggcggtggttccggaggcggcggctccggaggaggggggtcaagacaccctgacagtcaccaactcttcattggcaacctgcctcatgaagtggacaaatcagagcttaaagatttctttcaaagttatggaaacgtggtggagttgcgcattaacagtggtgggaaattacccaattttggttttgttgtgtttgatgattctgagcctgttcagaaagtccttagcaacaggcccatcatgttcagaggtgaggtccgtctgaatgtcgaagagaagaagactcgagctgccagggaaggcgaccgacgagataatcgccttcggggacctggaggccctcgaggtgggctgggtggtggaatgagaggccctccccgtggaggcatggtgcagaaaccaggatttggagtgggaagggggcttgcgccacggcagccagctggcgccctgatcaacgaggttgctgaacagctggtgggggtgtccgctgcttctgctgcggcctctgccttcgggtcattatccagcgcacttttgatgccaaaagatgggcgcactctggaggatgtggtgagagaactgctgcgtcctctgcttaaagaatggcttgaccaaaacctcccgaggatcgtcgagacaaaagtagaggaggaggtgcagcgcatctcccgcggacgcggcgcaTAACCTGGATGAGAGGCGTGTTAGAGAGTTTCATTAGCTTTAGGTTAACCACTGTTGTTCCTGATTGTACAAATACCAAGTGATTGTAGATATCTACGCGTAGAAAGTTAGGTCTAGTCCTAAGATCCGTGTAAATGGTTCCCAGGGAAGTTTTATGTACTAGCCTAGTCAGCAGGCCGCACGGATTCCAGTGCATATCTTAGTGATACTCCAGTTAACTCTATACTTTCCCTGCAATACGCTATTCGCCTTAGATGTATCTGGGTGGCTGCTCCACTAAAGCCCGGGAATATGCAACCAGTTACATTTGAGGCCATTTGGGCTTAAGCGTATTCCATGGAAAGTTATCGTCCCACATTTCGGAAATTATATTCCGAGCCAGCAAGAAAATCTTCTCTGTTACAATTTGACATAGCTAAAAACTGTACTAATCAAAATGAAAAATGTTTCTCTTGGGCGTAATCTCATACAATGATTACCCTTAAAGATCGAACATTTAAACAATAATATTTGATATGATATTTTCAATTTCTATGCTATGCCAAAGTGTCTGACATAATCAAACATTTGCGCATTCTTTGACCAAGAATAGTCAGCAAATTGTATTTTCAATCAATGCAGACCATTTGTTTCAGATTCGGAGATTTTTTGCTGCCAAACGGAATAACTATCATAGCTCACATTCTATTTACATCACTAAGAAGAGCATTGCAATCTGTTAGGCCTCAAGTTTAATTTTAAAATGCTGCACCTTTGATGTTGTCTCTTTAAGCTTTGTATTTTTAATTACGAAAATATATAAGAACTACTCTACTCGGGTAAATTGTGACTAACTAC
